# Effect of ligand sensing on flagellar bundle formation in bacteria

**DOI:** 10.1101/2021.02.11.430483

**Authors:** Megha Agrawal, Soumyadeep Chakraborty, Mahesh S. Tirumkudulu, K.V Venkatesh

## Abstract

*E. coli* swims in liquid media by rotating long appendages called flagella. The direction of rotation of each flagellum is governed by a transmembrane rotary nanomotor, which receives signals from ligand-specific receptors. Attractants bias the motor to rotate in CCW direction causing flagella to bundle and provide thrust for locomotion. Recent studies have shown that sensing not only leads to increase in CCW bias but also increases the motor rotation speed due to the recruitment of additional stator units bound to the rotor. Despite the detailed studies on bacterial motility, the effect of ligand sensing on the synchronization of flagellar filaments leading to bundle formation and changes in bundle geometry are not clear. In this work, we performed real-time imaging of the flagellar bundle of swimming cells in metabolising (glucose) and non-metabolisable (2-Deoxy-d-glucose) attractants. We characterized bundles during swimming by measuring visible distal length and the spread of filaments at poles. We show that sensing of attractant by receptor leads to the formation of tight bundles when compared to control buffer. Contrary to previous studies, the swimming speeds were proportional to the bundle tightness with the latter dependent not only on the bias but also on the torque exerted by the motor. We further show that the observed wiggles in the swimming trajectory of cells is directly proportional to the spread angles of bundle and is effected by both motor CCW bias and torque. Mutant cells, which were rendered non-motile due to the absence of the PTS (phosphotransferase system) sugar uptake mechanism, exhibited motility when exposed to the non-metabolisable attractant confirming that mere sensing can induce torque in flagellar motor. These results clarify the role of sensing and metabolism on bundle formation and its impact on the motility of cells.

**Statement of significance:** Peritrichously flagellated *E. coli* swims away or towards ligands by biasing the direction of rotation of its flagellar motor. Recently, it has been shown that motor speed is also modulated on merely sensing a ligand. How does this impact flagellar bundle formation and swimming behavior? Using real-time imaging, we show that the bundle geometry changes in response to both metabolisable and non-metabolisable ligand. Mere sensing of a ligand temporarily increases the motor torque and CCW bias that causes tight flagellar bundles and leads to smooth swimming trajectories at high speeds. Our result provides strong evidence of a new signalling pathway that controls the flagellar motor speed to enable the bacteria to respond efficiently to changes in its environment.

## Introduction

Bacteria employ a sensory system to explore their environment to find the best-suited niche for their survival. *E. coli* is one of the most studied bacteria for its sensing mechanism. It is rod-shaped and propelled by several thin helical flagellar appendages (about 4-7) emerging from random locations on their cell body. Each flagellum has a basal body, a flexible hook and a extended flagellar filament (≈ 5 *μ*m) [1] driven by a membrane-embedded, torque generating motor (≈ 50 nm) which rotates in either clockwise (CW) or counterclockwise (CCW) direction [2]. When a cell swims in a medium, each flagellar filament’s compliant hook enables them to self-organize into a coordinated bundle and push the cell in a forward direction. If one or more filaments change direction to CW, the flagellar filaments come apart from the bundle, causing a change in the swimming direction, termed as a tumble. Filaments are intrinsically left-handed helices but upon reversal to CW rotation, turn into right-handed (semi-coiled) with smaller diameter and pitch, that leads to dispersal of out-of-phase filament from the bundle [3]. Thus, tumbling occurs due to a combination of motor reversal and polymorphic changes in flagella and results in directional reorientation of the cell. When the cell swims, no external torque acts on the body and so the cell body rotates in a direction opposite to the rotating bundle so as to counter the torque exerted by the rotating bundle [4]. Three-dimensional tracking of *E. coli* reveals that the cell motion is akin to a random-walk motion constituting a sequence of runs (few seconds) and tumbles (fraction of a second) [5]. The directed motion towards attractants or away from repellents is thus achieved by a combination of run and tumble events. The swimming trajectory is not a straight line, but a helical curve of small radius [6], which is caused by the off-axis thrust exerted by sections of helical bundle. Large radius helical trajectories have also been observed in some bacteria due to either the off-axis position of the flagellar bundle relative to the cell body [7], or imbalance of mass distribution in the head [8].

In *E. coli*, a well-defined signaling pathway controls the direction of rotation of each flagellar motor. Five chemoreceptors specific for different ligands are clustered near the poles and they mediate signals via a cascade of proteins to the motor regulator protein, CheY [9, 10]. In the phosphorylated state, CheY-P binds to FliM component of the motor switch complex and enables it to rotate in CW direction [11]. The presence of attractants reduces the level of CheY-P; thus, the motor rotates in CCW direction in the null state [9] (Figure 1). Recent evidence suggests that sensing not only modulates directional bias but also increased swimming speeds by as much as 28% (23.3 *μ*m/s to 29.8 *μ*m/s) in *E. coli* in response to *α*-methyl aspartate gradients [12]. Run speeds were higher in metabolisable ligands such as glucose compared to its non-metabolisable analogue 2-Deoxy-d-glucose which are sensed by Trg receptor [13–15]. It was later shown that the increased swimming speeds correlates with the increased head rotation rates [14]. Further, the increased run speeds are accompanied with smoother swimming trajectories and reduced cell body wiggles [5, 13, 16, 17]. Inside the cell, the structure of flagellar motor is dynamic as stator units assemble and disassemble in response to changes in load, proton motive force, thereby altering the motor speed [18–20]. More recently, the increased swimming speed has also been attributed to recruitment of stators upon sensing of an attractant, which is indicative of a new signalling pathway for motor control [21]. While the aforementioned studies have investigated the influence of ligands either on the signalling pathway, motor control or the final swimming motion, an important component responsible for bacteria’s motion is also the bundle geometry and it’s dynamics.

**Figure 1:**
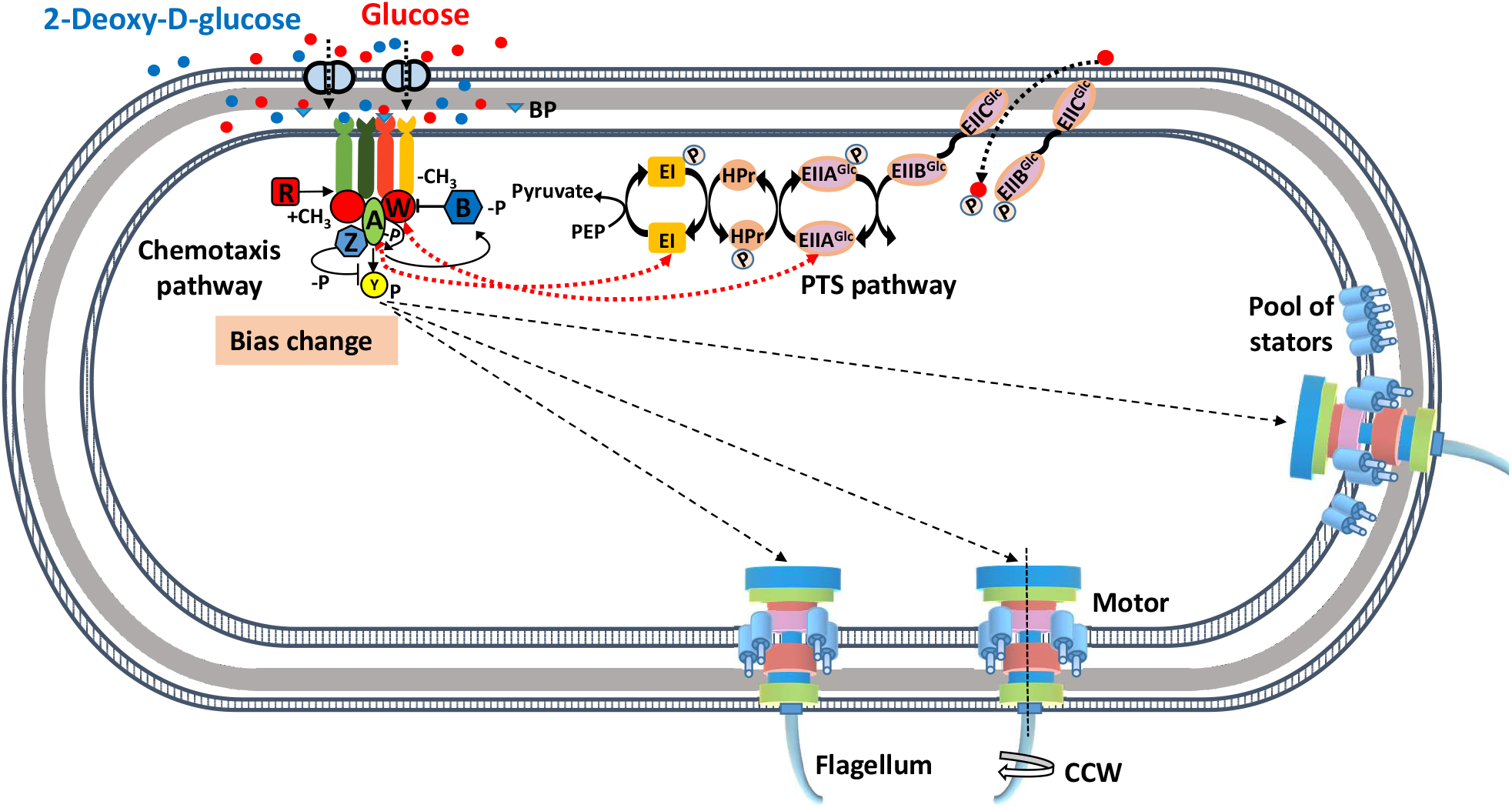
Transmembrane Trg receptor of the chemotaxis pathway signals the presence of attractants such as glucose and 2Dg to the flagellar motor via CheY protein. Phosphorylated state of CheY interacts with FliM of the rotor-switch complex of the motor, enabling them to rotate CW direction. Binding of an attractant activates CheA, which further activates the phosphatase activity of CheZ protein. CheZ dephosphorylates CheY and motor rotates uninterrupted in CCW direction. Additionally, glucose which is metabolisable ligand is also sensed by the PTS pathway whose signals integrate into the chemotaxis pathway. EI and EII both interact with CheA and CheW, and dictates the rotation via CheY [36, 37]. Such varied signals dictate the direction of rotation of individual flagellum placed at different locations of the cell. Sensing might synchronize these individual flagella into stable bundle formation by decreasing the probability of CW reversals and thereby impacting the swimming speed of cells. The above figure is adapted from Naaz et.al [21].

Much of the work on flagellar bundle has focused on the dynamics of bundle formation, via theory [22, 23], experiments [16, 24–26] and computer simulations [27–31]. In *E. coli*, the rotation of multiple flagella about their respective axis adds to the complexity in bacterial motion because rotational direction and speed are under the independent control of their motors. Interestingly, a cell with multiple flagella does not enable the cell to swim faster than a cell with a single flagellum since the thrust is a weak function of flagellar diameter [16]. Further, the tumbling probability is also independent of the number of flagella [32]. On the contrary, some marine bacteria with a single flagellum are able to choose a new swimming direction via buckling instability of the flagellum, also termed as a ‘reverse and flick’ mechanism [33] suggesting that ‘run-and-turn’ motion is possible even in the presence of a single flagellum. Soil bacteria such as *Rhizobium meliloti* are peritrichous like *E. coli* but their filaments undergo limited polymorphic transitions and are unidirectional as their motor can rotate only in CW direction [29]. Here, the bacteria change their swimming direction via rotational torque induced by applying different rotation speeds to individual flagellum. Thus it appears, that the role of multiple flagella in bacteria such as *E. coli* is to enable cells to choose a new direction after a tumble, *more efficiently*.

In the present study, we build on this understanding of bundle dynamics to investigate the influence of ligand sensing on the bundle geometry and the resulting swimming trajectory. Specifically, we determine how the sensing of metabolisable and non-metabolisable ligands influences the geometry of the flagellar bundle. The flagella geometry was visualized by tagging the flagellin proteins using amino-specific Alexa-fluor dyes, a technique pioneered by Berg and co-workers [34]. Since the compactness of the bundle is determined by both the motor bias and torque, we correlate the bundle geometry with motor performance via tethered cell assays. Cells were exposed to glucose, and its non-metabolisable analog 2-Deoxy-d-glucose (2Dg), which are sensed by the Trg receptor [10, 35]. Glucose is also sensed and metabolised by the phosphotransferase system (PTS) pathway [36]. Experiments were conducted with wild-type *E. coli* RP437 (WT) and its *trg*, *cheY*, *ptsI* gene deletion mutants. We show that both metabolism and sensing influence the bundle geometry via modulation of motor speed and bias, thereby effecting the motility of cell.

## Results

### Attractant sensing leads to smoother trajectories of *E. coli*

The wild-type *E. coli* cell population was exposed to a constant ligand concentration environment to measure the effect of sensing on the wiggling trajectories. The varying angular displacement about the mean swimming direction was quantified by measuring the apparent rotational diffusivity (D_*r*_) (Figure 2A, see Materials and methods section) of at least 3000 cells for each case. An increase in D_*r*_ signifies an increase in wiggling while a reduction pertains to smoother runs. Figure 2B presents the rotational diffusivity measured as a function of time for cells after they were exposed to 1000 *μ*M 2Dg and glucose along with the control (MB). Measurements were made at four different time points, namely, 1.5, 6.5, 11.5 and 16.5 min, from the start of the experiment. Figure 2B shows that the sensing of both attractants at two initial time points (1.5 and 6.5 min) resulted in 34-38% lower values compared to respective MB data points. Glucose had a more pronounced effect than 2Dg as the D_*r*_ gradually reduced to 55% and 74% of MB at 11.5 and 16.5 min, respectively. In the case of 2Dg, D_*r*_ remained 48% lower than MB at 16.5 min. All data points of glucose and 2Dg are significantly different from MB at *p* < 0.05 as computed by paired student t-test (except for 2Dg at 11.5 min). The trajectory of different cells in each buffer was also analyzed and the wiggling trajectories were approximated as helical paths in terms of pitch and radius (Figure 3B). In MB, the trajectory had a mean pitch of 1.50 ± 0.32 *μ*m and a mean radius of 0.47 ± 0.10 *μ*m (pitch/radius ± standard deviation) averaged over 20 cells. In 2Dg, cells showed smoother trajectories with a pitch of 0.88 ± 0.34 *μ*m and radius of 0.38 ± 0.05 *μ*m averaged from 20 cells (Figure 3B). The trajectories in uniform glucose were smooth with no detectable wiggling. These results clearly demonstrate that the sensing of attractants reduces the wiggling in swimming trajectory and the reduction is more significant in metabolisable ligand compared to its non-metabolisable analogue. The results are in agreement with earlier reported study [13].

**Figure 2:**
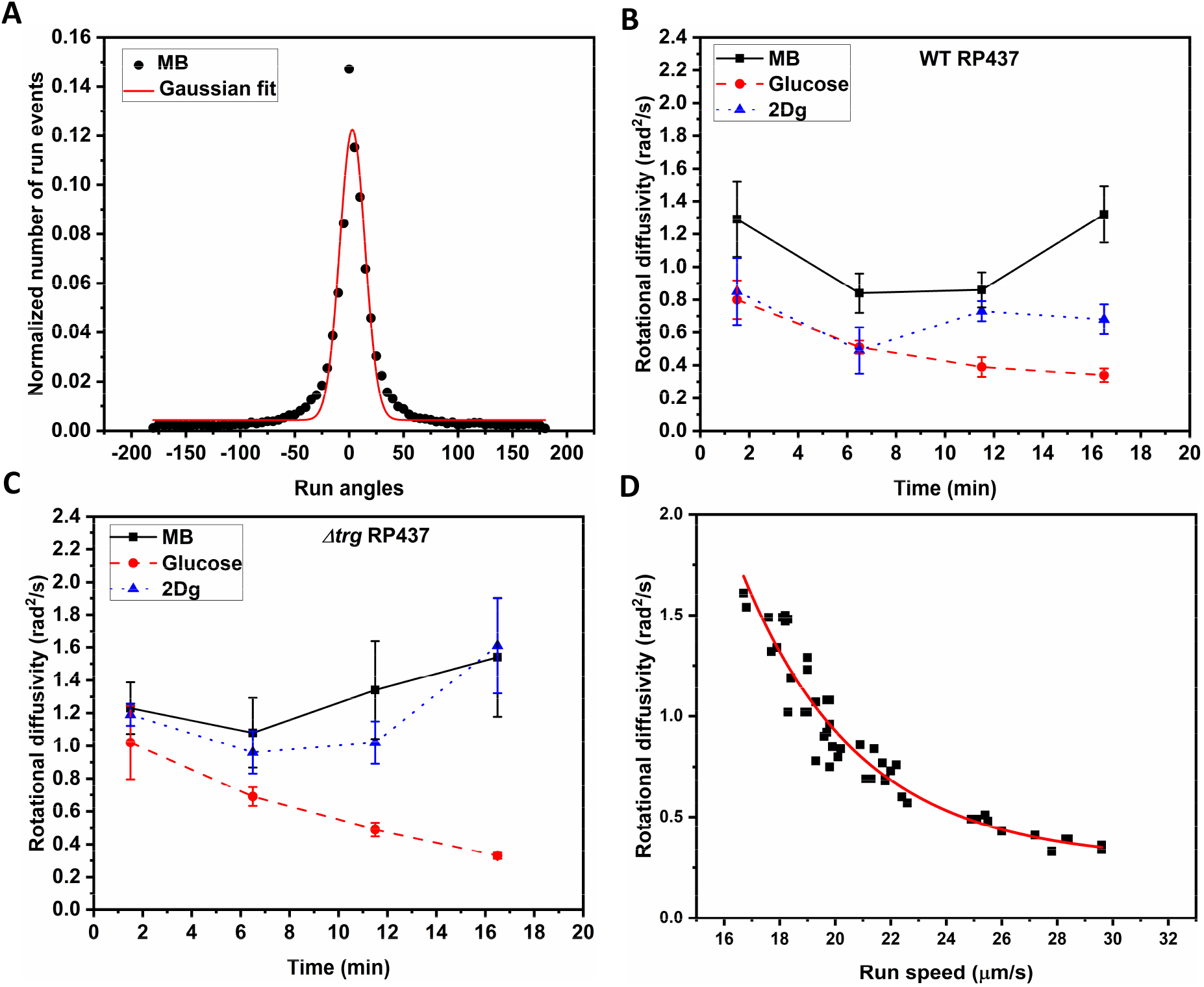
(A) The distribution of run angles for WT cell in MB (at 6.5 min) plotted against normalized number of run events for at least 3000 cells. The Gaussian curve is fit to obtain the standard deviation 〈*θ*〉^2^ so as to calculate the *apparent* rotational diffusivity, 〈*θ*〉^2^/2*t* where *t* = 0.047s. Variation in rotational diffusivity with time in 1000 *μ*M glucose and 2Dg as compared with control buffer (MB) for (B) Wild type RP437. All data points of glucose and 2Dg are significantly different from MB at *p* < 0.05 as computed by paired student t-test (except for 2Dg at 11.5 min). (C) Δ*trg* RP437. All data points of glucose are significantly different from MB at *p* < 0.01 as computed by paired student t-test. The difference in values of MB and 2Dg were insignificantly different from each other (*p* > 0.05.) except at 11.5 min. The time axis represents incubation period after ligand introduction. Error bars are standard errors from four independent experiments. (D) Rotational diffusivity is inversely proportional to the run speed of cells. Swimming speed and rotational diffusivity of each WT, Δ*trg*, Δ*ptsI*, and Δ*cheY* RP437 strains were measured in MB, glucose, and 2Dg at four time points, 1.5, 6.5, 11.5, and 16.5 min. Each data point is an average of at least 2500 cells from four independent experiments. The solid curve is an exponential fit for the data points.

**Figure 3:**
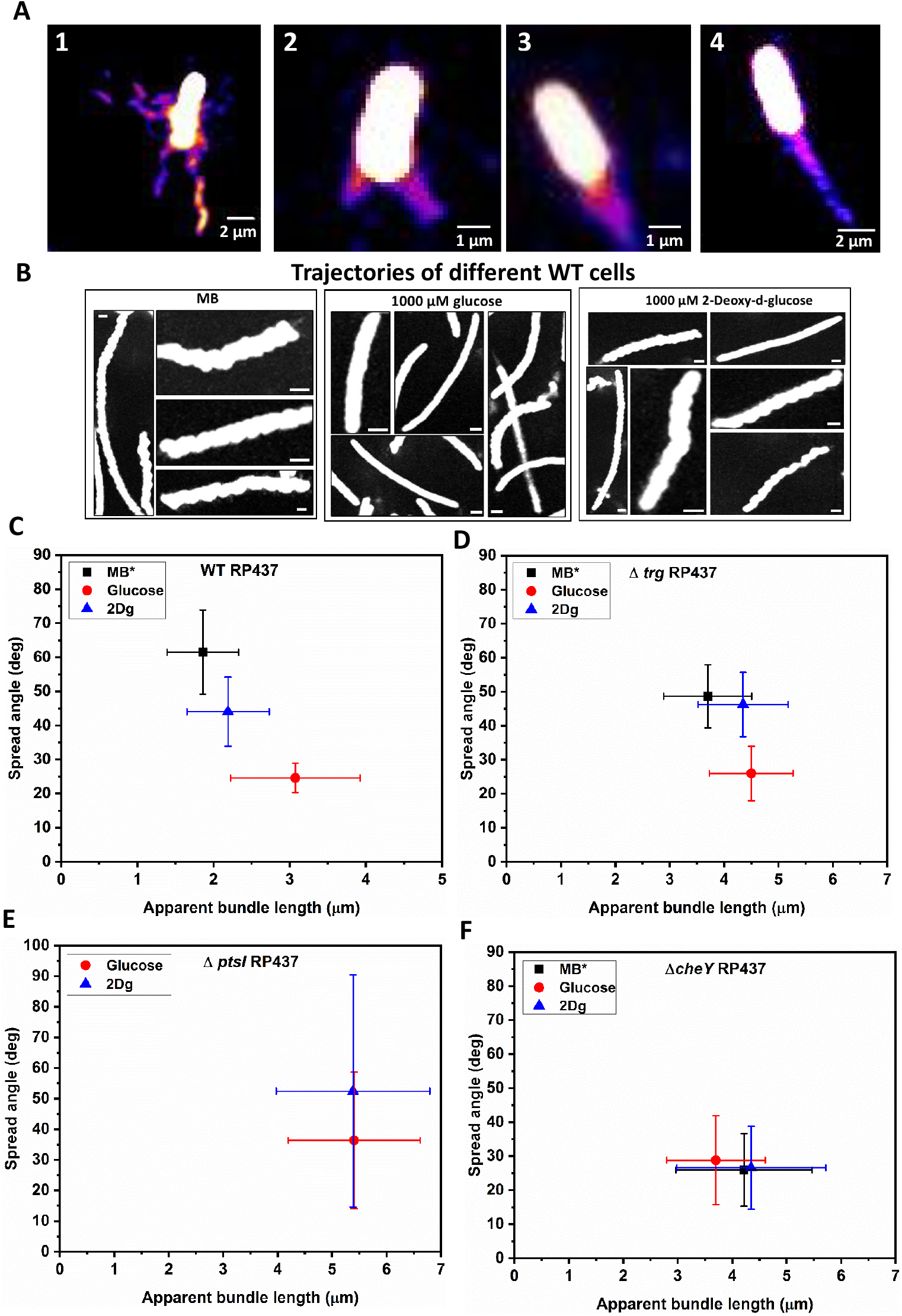
(A) Different configurations of flagellar bundle in WT 1) Stationary cell stuck to the glass showing four flagella. Swimming WT cells in 2) MB* with two flagellar bundles 3) 2Dg shows wider bundle spread at poles 4) Glucose shows tighter bundles at poles. (B) Trajectories of different WT cells in motility buffer shows wiggling while in glucose smooth trajectories were seen. Cells in 2Dg showed both wiggling as well as smooth trajectories (Scale bar, 2 *μm*). (C) To characterise bundle geometry, spread angles and length of the flagellar bundle were measured in MB*, 1000 *μ*M glucose, and 2Dg for WT. The spread angles in different ligands were significantly different from MB* at *p* < 0.001 as computed by one-way ANNOVA. (D) In Δ*trg* RP437, the spread angle values in glucose were significantly different from MB* at *p* < 0.001 as computed by one-way ANNOVA (E) In Δ*ptsI* RP437, no data is shown for MB* since cells had no net motion with unbundled flagella. The spread angles in 2Dg was not significantly different from glucose at *p* > 0.05 as computed by one-way ANNOVA (F) Δ*cheY* RP437. Each data point is an average of 25-40 cells. The data was averaged from three independent cell cultures.

To further confirm that reduction in the D_*r*_ values is indeed an outcome of ligand sensing via the transmembrane receptor, experiments were repeated with *trg* gene mutant strain. Figure 2C shows that there was no significant difference in the values of D_*r*_ between MB and 2Dg. The initial reduction in case of glucose was only 17% of MB (0 min), whereas in WT, it was 38%. However, glucose is also sensed by the phosphotransferase uptake pathway whose signal is integrated with the chemotaxis pathway [36]. Therefore, with time the D_*r*_ values gradually reduced to 78% of MB at 15 min, similar to that achieved in WT.

### Swimming speed is inversely related to rotational diffusivity

To find the relation between swimming speed and rotational diffusivity, both parameters were measured for WT and its mutants (Δ*trg*, Δ*cheY*, Δ*ptsI* RP437) in glucose and 2Dg at different times after ligand exposure. Figure 2D shows the consolidated graph where each data point is an average of at least 3000 cells. The speed of the cell population varies inversely with respective D_*r*_, as depicted by the fitted-solid curve.

### Glucose metabolism and sensing leads to tighter bundles

To explore the reason for reduced angular displacements during sensing, flagellar proteins were stained to visualize the flagellar bundle of cells while swimming. The fluorescent-labeled WT cells were exposed to control buffer (MB*, which has a different pH value as compared to MB, see Materials and Methods section), and 1000 *μ*M of glucose and 2Dg. Figure 3A shows different configurations of flagellar bundles observed when a batch of WT cells was imaged in the uniform concentration of MB*. An average of 3-4 flagella were observed with curly configuration for a stationary cell (Figure 3A-1). Figure 3A also shows bundles of swimming cells formed by WT in different ligands as compared to control buffer. Two bundles are seen in MB* (Figure 3A-2). The bundle in 2Dg shows a wider spread at the pole (Figure 3A-3) while glucose has a smaller spread angle (Figure 3A-4). The flagellar bundle in glucose appears as a long blurred tail which is in accordance to previous study [38]. However, neither the helicity of individual flagellum nor the total number of flagella in an *E. coli* could not be detected due to the high rotation speed of the flagella.

For quantifying bundle geometry, its length and spread at the pole were measured as described in the supplementary information (see supplementary information, S1A). The spread angle quantifies the compactness of the bundle. The bundle length reported here is the measured projected length and may not be the true measure of flagella length since flagella length is highly dependent on culture condition, cell preparation methodology and imaging condition. The apparent bundle lengths across various mutants in different ligands were observed to be in the range of 1.5-6 *μm* which is in agreement with the values reported earlier in Alexa fluor 488 labelled *E. coli* AW405 (5.6 ± 2.9 *μm*) [24]. Also, the filaments lengths were shorter in Alexa fluor 488 as compared to other dyes due to its chemical effects that might render the filaments brittle [24]. Since the goal is to determine relative changes in bundle length and spread angle, the shortness of individual filaments is not expected to influence the final results. Images of about 25-40 individual cells, each of WT, Δ*trg*, and Δ*ptsI* RP437 strains, were analyzed to extract these two parameters, each in MB*, glucose, and 2Dg environment (Figure 3C-F). WT showed an average spread angle of 62°±12° (angle ± standard deviation) in MB*, an intermediate spread of 44°±10° in 2Dg and the smallest spread angle of 24°±4°in glucose (Figure 3C). In MB*, cells exhibited either loose single bundles or multiple bundles. On the other hand cells exposed to 2Dg and glucose showed only a single bundle. Apparent bundle length varied inversely with spread angle with the longest bundles observed in glucose. Thus, the sensing of ligand appears to synchronize the movement of individual filaments so as to form long, single, tight bundles.

A similar experiment was also conducted with Δ*trg* RP437. It showed an average flagellar spread angle of 48°±9.2° in MB* and 46°±9.4° in 2Dg, whereas in glucose it was around 26°±7.9° (Figure 3D). The absence of Trg receptor eliminates signal transduction due to sensing of 2Dg leading to spread angles similar to those observed in MB*. These results confirm the analogous trend observed for D_*r*_ (Figure 2C). The bundling parameters in glucose were unaffected by Trg receptor’s absence and were similar to those observed for WT in glucose. It is evident from these experiments that sensing of glucose by Trg receptor and PTS uptake system both result in tighter flagellar bundles. This is not surprising given that the PTS uptake system reduces the phosphorylation of CheA, which in turn decreases the concentration of CheY-P, and therefore biases the motor towards CCW rotation.

Since glucose metabolism is controlled by the PTS pathway, Δ*ptsI* RP437 strain was used to investigate the role of sensing via PTS pathway on the compactness of bundles. Δ*ptsI* cells showed an increased spread angle of 36°±22° in glucose, which is highest compared to WT and Δ*trg* strains in the same media. The spread angle for 2Dg was 52°±38° (Figure 3E). We could not obtain sufficient data in MB* since flagella were completely unbundled leading to negligible motility.

### Absence of CheY response regulator resulted in tight bundles

The response regulator protein, CheY was deleted, and the flagellar bundle parameters were measured. The responses for MB*, glucose, and 2Dg were indistinguishable as all cells showed tight bundles (Figure 3F). The spread angle were in the range of 26-30 °(±12°). The values of the spread angle were similar to that of glucose as obtained for WT and Δ*trg* cells. This suggests that when all the flagella rotate in the CCW direction, the bundle is compact leading to negligible wiggling.

### Swimming speed is inversely related to spread angle

Since ligand sensing synchronizes the flagellar rotation to form tighter bundles in presence of ligands as compared to control buffer, we investigated whether this influences the run speed of cells. The run speed of WT cells in MB*, glucose, and 2Dg were plotted against their respective average spread angle. Figure 4A shows that speed was inversely related to the spread angle as depicted by the exponential fit. Tight bundles with the highest speeds were obtained in glucose reaching 23 *μ*m/s. For 2Dg, all data points lie in between those for glucose and MB*. The maximum speed was around 13 *μ*m/s in 2Dg. In MB*, we obtained considerably lower speeds as compared to earlier reported speeds of 18.8 ±8.2 *μ*m/s [39]. The reduced speed for the dyed cells can be attributed to not only shorter bundles but also to the effect of the laser beam. It is well known that light of wavelength 510 nm or lower reduce the the motility of bacteria [38]. Nevertheless, these factors do not effect the conclusions as we draw all comparisons for glucose and 2Dg with respect to the control buffer subject to identical experimental conditions. Cells in MB* exhibited mostly irregular bundle formation (including multiple bundles), and thus, the maximum speed was about 9 *μ*m/s, which is much lower as compared to glucose and 2Dg. The measured run speed of cells was significantly different in MB*, glucose, and 2Dg with *p* < 0.01 as computed by one-way ANNOVA. The apparent bundle length was inversely related to the spread angles (Figure 4B) suggesting that thick bundles are visible more easily.

**Figure 4:**
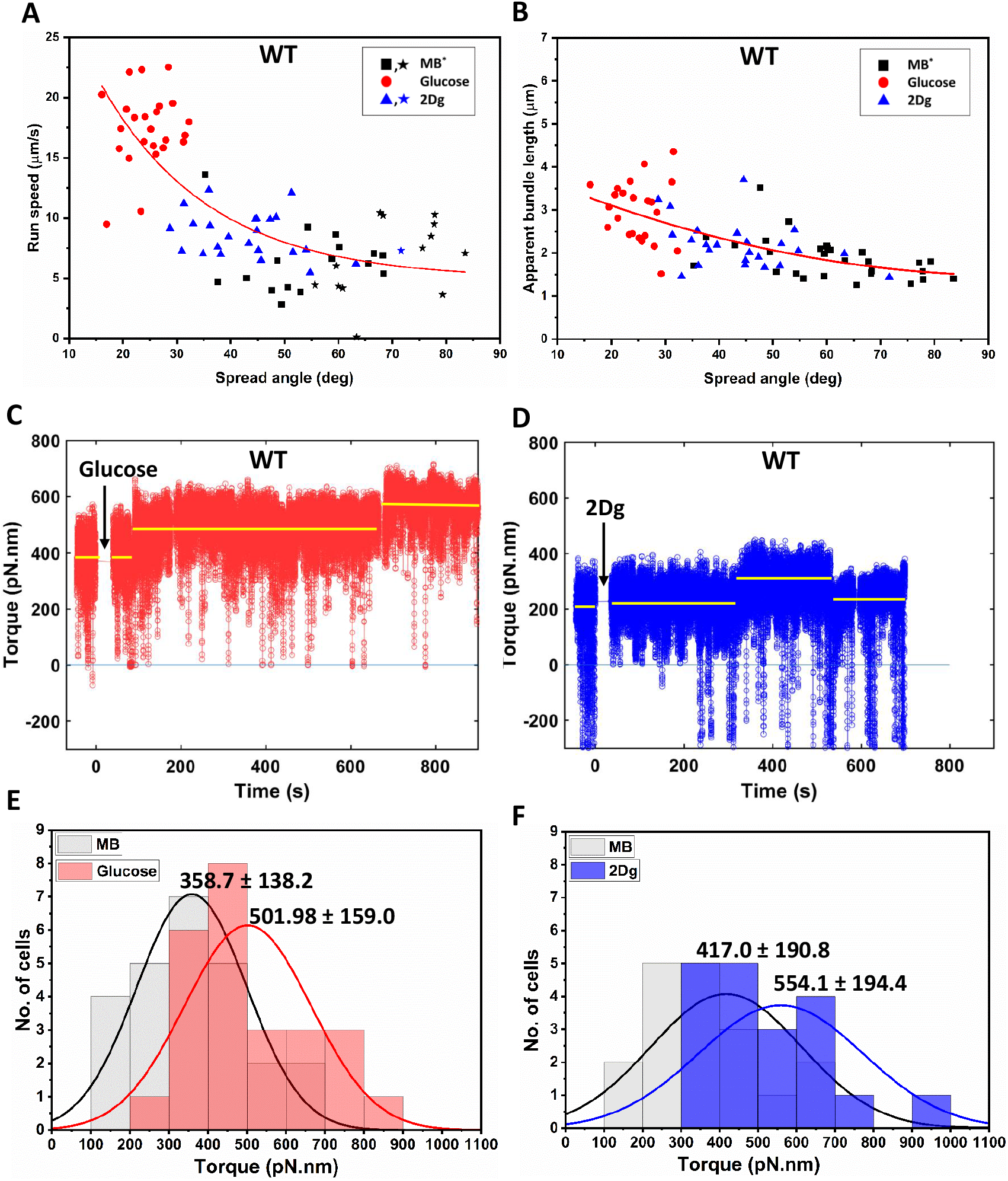
(A) The spread angle attained by WT cell while swimming in MB*, glucose, or 2DG is plotted against its respective run speeds. Cells with higher swimming speeds tend to have the lowest spread angle as evident for glucose, while for 2Dg, values lie in between MB* and glucose. Speeds in each ligand were significantly different from each other at *p* < 0.05 as computed by paired student t-test. Star marked symbol denotes cells having improper bundling, i.e multiple flagellar bundles or unbundled flagellar filaments. (B) Bundle length of each WT cell plotted against its spread angle. Tethered cell experiment with WT cells were performed to determine changes in motor torque initially incubating in MB and exposed to 1000 *μ*M glucose or 1000 *μ*M 2Dg. A single motor (WT) showed increase in motor torque when suddenly exposed to (C) 1000 *μ*M Glucose (arrows indicate time of introduction). The response was sustained for 900 seconds. (D) 1000 *μ*M 2Dg but the response was limited to 500 seconds. The torque values depicted are moving average of 0.1 s (10 frames). Similarly, the torque for more than 20 paired cells that were initially incubating in MB increased when exposed to (E) 1000 *μ*M glucose and (F) 1000 *μ*M 2Dg. Values for glucose and 2Dg are obtained after 16.5 min and 6.5 min of exposure, respectively. The population means and standard deviations are included in the figure. All values are significantly different from MB at *p* < 0.001 computed by paired t-test. Data was obtained from four independent experiments.

Swimming speeds were also measured in Δ*trg* RP437 cells and correlated with the spread angle of bundle. There was no difference in run speed and spread angles for cells in either MB* or 2Dg (See supplementary information, S4A). Average run speed in MB* and 2Dg was 14.5 ± 4.5 *μ*m/s (speed ± standard deviation) whereas in glucose it reached 22.8 ± 5.6 *μ*m/s. This result confirms that sensing via the Trg receptor and metabolism via the PTS uptake pathway, both lead to smaller spread angles. Further, the data presented in Figure 2D and Figure 4A show that the rotational diffusivity of a cell is directly correlated to the bundle spread angle.

### Tighter bundles are due to the combined effect of increased motor torque and CCW bias

Δ*cheY* RP437 was studied under ligand conditions similar to that for other strains (1000 *μ*M glucose and 2Dg) to probe the effect of increased CCW bias on bundle formation and motility of bacteria. Cells showed low spread angles of 26-29°±11° irrespective of ligand condition. However, run speeds in glucose and 2Dg were significantly different from MB at *p* < 0.05 as computed by ANNOVA one-way (Figure 5). These results suggest that an increased motor speed/torque in addition to a constant CCW bias may be responsible for these observations.

**Figure 5:**
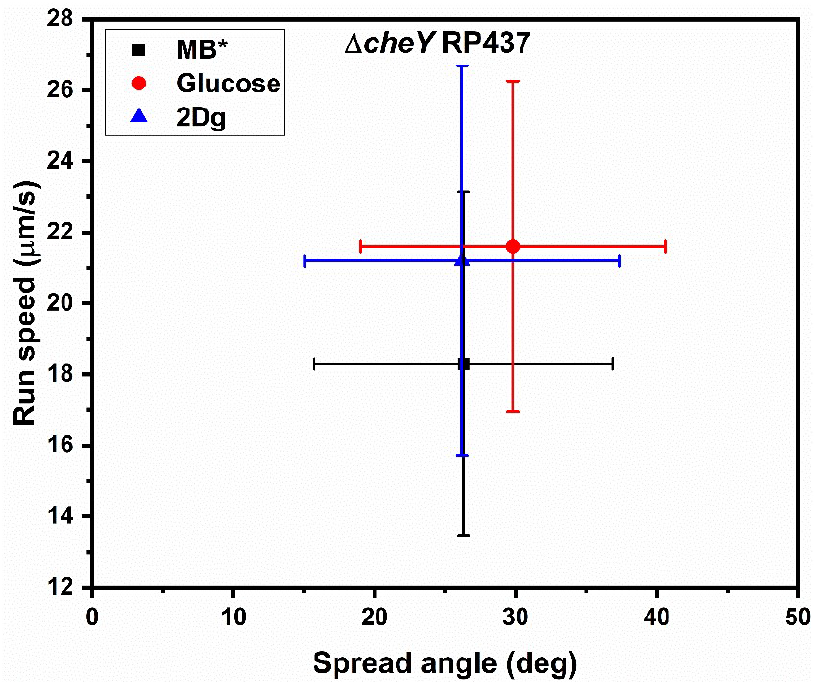
Variation of run speed in response to 1000 *μ*M glucose and 2Dg in Δ*cheY* RP437 cells. Each data point was an average of 30-35 individual cells. Error bars depict standard deviation. Run speeds in glucose and 2Dg were significantly different from MB at *p* < 0.05 as computed by ANNOVA one-way

We further investigated these observations by determining the motor torque for WT cells, where the cells were tethered on glass slide. We have shown recently that the motor speed increases when the cells sense a ligand and the mechanism is independent of chemotaxis pathway [21]. By measuring the rotational speed of the tethered cell using the same methodology and then calculating the motor torque (see supplementary information for details), we could determine the influence of the ligands on the torque output of individual motors. In WT, the motor torque increased from 380 pN.nm in MB to 475 pN.nm after 100 s and 570 pN.nm after 650 seconds of exposure to glucose (Figure 4C). Similarly, in response to 2Dg, motor torque increased in single step from about 200 pN.nm in MB to 300 pN.nm at around 300 seconds after exposure to 2Dg (Figure 4D). The increase in torque in response to glucose (40%) and 2Dg (33%) was confirmed by averaging over at least 20 cell pairs (Figure 4E-F).

## Discussion

The directional motion of bacteria is dependent on the stability of flagellar bundle which is subjected to intrinsic noise arising from individual flagellar motors. Owing to the nanometer dimension of flagellum and high rotation speed of flagellar bundle (≈ 100 Hz), it is a challenge to observe the cell body and the geometrical features of the flagellar bundle simultaneously. Hence, much of the previous understanding of the bundling process comes from either simulations or macroscopic scale models [25, 26]. The bundling has largely been explained by hydrodynamic interactions between flagellum present in a viscous fluid. It is a combination of helix handedness, motor torque and a sense of motor rotation [25, 26, 29, 40]. Motor reversals have been attributed to the binding of CheY-P to the motor protein FliM [11] while Berg and others [16, 24] have highlighted the morphological transformations of the flagella during bundling and unbundling. However, what has not been investigated is the geometry of flagellar bundle in the presence of ligands, both metabolisable and non-metabolisable, and how these changes influence the swimming trajectory of bacteria. Thus real-time experiments linking changes in bundle geometry due to ligand sensing to the swimming trajectory are largely missing.

To this end, we chose attractants, glucose and its non-metabolisable analogue, 2Dg, which are both sensed by the Trg receptor. Note that glucose is a preferred carbon source for *E. coli* and hence is an appropriate ligand to investigate its influence on bundle geometry and the swimming trajectory. We performed a population-level study of WT strain to measure the change in rotational diffusivity in presence of ligands as a function of time. Our experiments with WT strain in glucose showed that the rotational diffusivity (D_*r*_) reduced with increasing time, an indication that there is reduced wiggle in its swimming trajectory. Even for the non-metabolisable ligand, D_*r*_ was decreased by about 35% (compared to that in MB) for a short time after exposure to the ligand (Figure 2B). The absence of a response to 2Dg for the Δ*trg* RP437 strain clearly showed that sensing via receptor influenced the motor motion in WT cells so as to reduce their rotational diffusivity leading to smoother runs. These results were confirmed via control experiments with Trg receptor-deficient strain, where no such change in swimming trajectory was observed upon sensing of 2Dg. However, this was not the case for the Δ*trg* RP437 strain exposed to glucose. The reduction in rotational diffusivity was similar to that observed in WT, suggesting that the metabolism of glucose via the PTS pathway could also lead to smooth runs (Figure 2B). The inverse relation between D_*r*_ and swimming speed was evident from consolidated population data of WT and its mutant strains in different ligands (Figure 2D).

How do cells achieve differential swimming speeds? Does the answer lie in geometrical features of the bundle, which is ultimately responsible for cell propulsion? A flagellum rotates stochastically in either CW or CCW direction. The body rotates in opposite direction to balance the torque generated by rotating bundle, therefore interactions between individual flagella becomes important. A stable bundle cannot form if all the flagella change direction continuously which would otherwise lead to jamming. Although it is well known that sensing of ligand increases the CCW bias of flagellar rotation, it is not clear as to how it effects the bundle formation. We performed real-time imaging of *E. coli* WT cells in control motility buffer to visualize the bundle. Swimming cells with loose bundles and in some cases, multiple bundles, were observed. The latter are a common occurrence in multi-flagellated bacteria such as *Bacillus subtilis* [41] although it has not been reported previously for *E.coli*.

We quantified the configuration of bundles by measuring two parameters, namely, the spread of filaments around the pole and apparent bundle length. In WT, the majority of cells in MB* had irregular bundles (Figure 3C). Tight bundles (lowest spread angle) were observed for glucose sensing by WT and, *trg* and *cheY* mutant strains (Figure 3C-D, F). However, it was not the case for *ptsI* mutant (Figure 3E) indicating that combined effect of sensing and metabolism results in tight bundles in glucose.

The spread angles for WT in 2Dg were intermediate to those observed in MB* and glucose (Figure 3C). Further, the spread angles matched with those obtained in MB* for the Trg-receptor deficient strain (Figure 3C). We then asked as to what would happen to the flagellar bundle if all motors rotate exclusively in CCW direction. In this regard, experiments were performed with Δ*cheY* RP437 [42] and in all three cases (MB*, glucose, 2Dg), the bundles were tight with low spread angles (Figure 3F). In fact, the spread angle values were similar to those obtained for glucose in WT.

Does tight bundle confer any advantage to a swimming cell? Our study shows that cells respond to attractants by forming tight bundles. Cells with improper bundles (loosely stacked filaments or multiple bundles) result in the slowest run speeds (Figure 4A). Interestingly, when PTS was absent, there was no significant motion in MB* but cells exhibited directed motion when exposed to 2Dg. This is remarkable because it shows that mere sensing could induce non-motile cells to swim, which goes beyond the current understanding of sensing only influencing the bias of flagellar motor.

A consolidated data of all cells with their spread angles shows an inverse relationship with their corresponding run speeds (see supplementary information, S4B). These results appear contrary to earlier reports [16, 17] where it was reported that swimming speeds were not the outcomes of tight or loose bundles [17]. The aforementioned experiments were performed in the presence of glucose [16] and in the presence of methyl cellulose, both of which result in tight bundles and hence it is not possible to obtain a large variability in bundle geometry. While our results clearly show that tight bundles are achieved due to CCW bias, one of the most important results of the study is that swimming speed can be modulated separately via sensing and metabolism. Experiments with Δ*cheY* mutant showed higher swimming speeds in glucose and 2Dg as compared to MB*, even though the motor was rotating continuously in CCW bias (Figure 5).

Similarly, the D_*r*_ values for Δ*cheY* were reduced in glucose (57%) and 2Dg (43%) as compared to MB, confirming that wiggles were further reduced during sensing even though the cells were completely biased to rotate in CCW direction (see supplementary information, S3). Further confirmation of these results was obtained from tethered cell studies with WT which revealed that the motor torques also increased due to sensing (Figure 4C-F). The reported values of torque in MB for WT are in agreement with those reported by Darnton et. al [16].

One of the most important observation from this work is that mere sensing of a ligand temporarily increases the motor torque and CCW bias that causes tight flagellar bundles and leads to smooth swimming trajectories with high speeds. This effect was observed even in Δ*cheY* mutant strain suggesting that chemotactic pathway is not responsible for the increase in motor speed. Knocking-off the sensor eliminates this effect while the motility of Δ*ptsI* mutant cells could be recovered via mere sensing of a ligand. These results clearly indicate the existence of a hitherto unknown signalling pathway that connects the sensor to the motor. In the presence of glucose, where the ligand is both sensed and metabolised, the bundles were the tightest and the highest swimming speeds were achieved. Clearly, *E. coli* has evolved a complex signalling system that couples to a sophisticated motor and enables the bacteria to respond in varying degrees to changes in its environment.

## Materials and Methods

### Bacterial strains and growth condition

All mutants are derivatives of *E. coli* RP437 strain as described in Table 1. Mutant strains Δ*trg* and Δ*ptsI* were developed by following one-step gene inactivation method [43, 44] and the details are reported elsewhere [21]. Δ*cheY* strain was a generous gift from Prof. J. S. Parkinson. Cells were streaked from glycerol stock into tryptone agar plates. A single colony isolate was grown overnight and then sub-cultured in tryptone media (0.015g/l; Sigma) till mid-log phase (OD _600_ at 200 rpm, 30°C). All experiments were performed at room temperature 23°C.

**Table 1:** Bacterial strains used in this study

| Strain | Relevant genotype | Parent strain | Comments, references |
| --- | --- | --- | --- |
| *E. coli* RP437 |  |  | *E. coli* wild type [42] |
| RP5232 | $\Delta cheY$ | RP437 | [42] |
| MTKV01 | $\Delta trg::Frt$ | RP437 | [21] |
| MTKV04 | $\Delta ptsI::Frt$ | RP437 | [21] |

### Determination of rotational diffusivity and run speeds at the population level

Bacterial cell culture (50 ml) was centrifuged at 4000 rpm for 10 min to obtain a pellet. The pellet was washed twice via resuspension in motility buffer (MB). MB is composed of *K*_2_*HPO*_4_, 11.2 g; *KH*_2_*PO*_4_, 4.8 g; (*NH*_4_)_2_*SO*_4_, 2 g; *MgSO*_4_*.H*_2_*O*, 0.25 g; Polyvinylpyrrolidone, 1 g and EDTA, 0.029 g per liter of distilled water. Cells were introduced in vials containing MB, 1000 *μ*M glucose, and 1000 *μ*M 2Dg such that the final bacterial concentration was approximately 10^6^−10^7^ cells/mL. The cell suspension were introduced into rectangular glass microchannels of dimension, 5 cm (L) *×* 1000 *μ*m (W) *×* 100 *μ*m (H) (VitroCom Inc.) via capillary action. Both the ends of the microchannel were sealed with wax after the introduction of cells. For each sample, a new capillary was used at 0, 5, 10, 15 min to measure the change in rotational diffusivity with time. The measurements were done over 3 min.

An inverted microscope (IX71, Olympus) fitted with Evolution VF cooled Monochrome camera (Media cybernetics, Japan) was used for imaging. Trajectories of swimming bacteria were recorded with 40X (0.75 NA) objective using a dark-field condenser. At each time point, six videos each of about 15 s duration were recorded over a span of 3 min at a frame rate of 21 fps. Images were taken in the central region of the micro-channel away from the channel walls. The trajectories of the cells were obtained using commercial software, ImageProPlus. The data were analyzed using an in-house code written on MATLAB to obtain the rotational diffusivity from more than 2,500 cells for each condition. We considered cells within 1 *μ*m of the focal plane with tracks longer than 0.5 s, which ensured that all out-of-plane motions were ignored by the analysis. A tumble event was identified when the swimming speed of the cell was below half the mean swimming speed, and the change in the turn angle was higher than 4° between successive frames (at 21 fps) [45]. The measured average swimming speed of ± 7.9 *μ*m/s (average ± standard deviation) and an average turn angle of 71° for RP437 cells dispersed uniformly in a micro-channel containing plain motility buffer are close to those observed for the same strain reported earlier, 18.8 ± 8.2 *μ*m/s and 69° [39]. The rotational diffusivity is defined as D_*r*_ = 〈*θ*〉^2^/2*t*, where *θ* is the angular displacement between two consecutive frames, t is the time difference between those two frames (0.047 s) and the brackets represents a grouped average of *θ*. The details of the analysis have been described in earlier published reports [13, 14].

### Fluorescent staining of flagella

Bacterial culture of OD_600_ ≈ 0.6 (10 ml) was pelleted by centrifugation (4000 rpm, 10 min). Pellet was washed thrice by centrifugation (2000 rpm, 10 min) in pH 7.0 Buffer A (0.01 M KPO_4_; 0.067 M NaCl; 10^*−*4^ M EDTA; 0.001% Tween 40). After the last wash, cells were resuspended in 250 *μ*L Buffer B (same as Buffer A but adjusted to pH 7.5) for staining. 20 *μ*L of Alexa fluor 488 NHS ester (or succinimidyl ester) (ThermoFisher Scientific, product number-A20000) (5mg/ml) was added to cell suspension [34] and the cells were incubated in the dark at 30°C for one hour with gentle mixing at 100 rpm. Excess dye was removed by washing multiple times in Buffer A (2000 rpm, 10 min). Since maintaining a pH of 7.0 is critical for stable staining, buffer A was used for dissolving ligands. Hence, for dye experiments MB* refers to plain buffer A. A dilute suspension of cells was obtained by adding 1 or 2 *μ*l of cell pellet in each of 500 *μ*l MB*, glucose or 2Dg. Cell suspension was then introduced in oxygen permeable device and immediately subjected to imaging. It should be noted that buffer was not exchanged during the experiment and different devices were used for each ligand.

#### Preparation of slides and image acquisition

Rectangular chambers were formed by placing two parallel silicone rubber spacers on glass coverslip. Thin-sheets of PDMS (Polydimethylsiloxane) were placed on top of the spacers. This setting ensures permeability of oxygen to the cells and thus, does not affect the motility of cells due to the oxygen limitation during the course of our experiment.

The cell suspension was introduced in these channels. Imaging was done using spinning disc confocal microscopy (Perkin Elmer UltraView system with Olympus IX71) with 100X (1.4 NA) dark phase oil-immersion objective. Setup was excited with 488 nm wavelength laser and exposed for 46 ms. The system was set at 1-by-1 binning and 40% laser power. Images were recorded at a speed of 17 fps by an EMCCD camera (Hamamatsu Inc.). Exposure time for each frame was 46 ms. Since average bundle rotation rate is 131 Hz [16], this means that imaging essentially measured the bundle shape averaged over 5-6 flagellar bundle rotations.

#### Image Analysis and characterization of flagellar bundles

The cell body appeared very bright as compared to flagella. Hence, brightness and contrast were adjusted using ImageJ (NIH) software to visualize flagella distinctly using ImageJ. Individual swimming cells were cropped from the main movie. For each cell, three parameters were measured – spread angle, bundle length, and speed. Spread angle of a bundle at the pole was measured by forming an angle at a fixed distance of 1 *μ*m away from it such that the complete bundle is contained within this angle. Bundle length was measured as the distance between the polar end of the cell body and the distal end of the flagellar bundle (see supplementary information, S1A). For each cell, these two parameters were measured in at least three different frames, and an average was computed. The swimming/run speed of cell was measured by determining distance traveled between frames and was calculated using the following equation

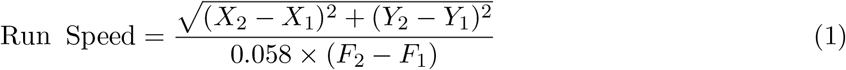

where *X*_1_, *Y*_1_ and *X*_2_, *Y*_2_ are coordinates of centroid of cell in frames F1, F2 respectively and 0.058 s is the time difference between consequent frames. For each strain and ligand condition, 25-40 cells were measured, which is similar to the number of cells used to determine bundle properties in earlier studies [16]. All distances were calibrated by recording images of an objective micrometer.

### Determination of motor torque by tethered cell experiment

Tethered cell experiments were conducted as described in detail previously [21]. Briefly, mid-log phase grown cells were sheared by passing through 21-gauge syringe needle (75 times) and anchored to glass-slide pre-incubated with anti-flagellin antibody (Figure 6). The imaging was done with an inverted microscope (Olympus IX71) fitted with 100X (1.4 NA), oil-immersion objective. Area was scanned for rotating cells and videos were captured at an acquisition rate of 60 fps for 30 to 44 s using a CMOS camera (Hamamatsu Inc). In order to measure the response of single flagellar motor to ligands, we first measured the response in MB and again after adding glucose/2Dg. Such pair-wise assessment removed any variability arising due to cell size and tethering geometry. The image analysis was done in ImageJ and motor speed calculation were made using in-house MATLAB code described previously [21]. The torque of each rotating cell based on their shape was then calculated as described previously [46] and is detailed in supplementary section.

**Figure 6:**
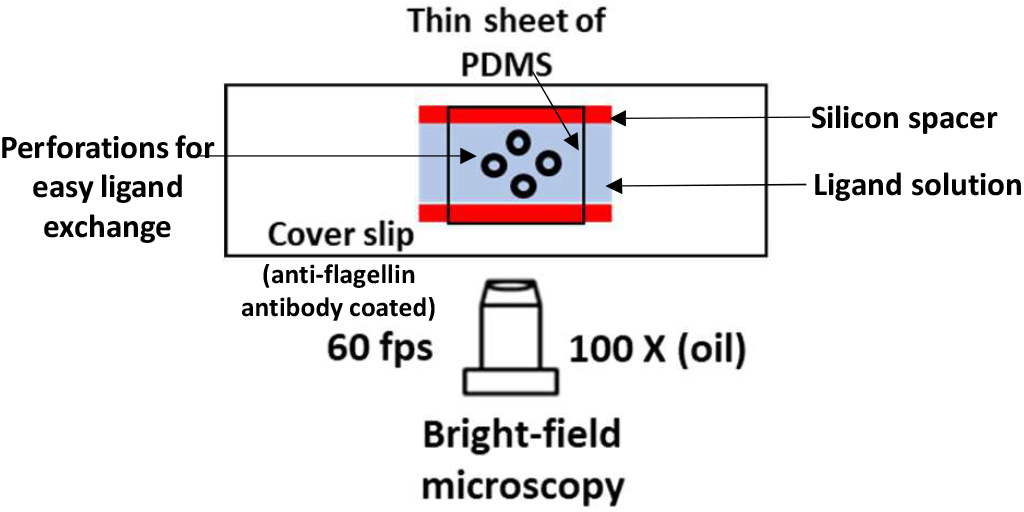
Schematic of setup for tethered cell experiment of *E. coli*. Cells were first imaged in MB, and then concentrated ligand solution was introduced through holes.

## Acknowledgments

We thank Prof. John S. Parkinson for providing us with *E. coli* RP437 wild-type and its *cheY* deletion mutant strains. Financial support from the Department of Science and Technology, India (SB/S3/CE/089/2013) and Department of Biotechnology, India (BT/PR7712/BRB/10/1229/2013) is acknowledged.

## Author contribution

M.A., S.C., M.S.T. and K.V.V. designed research; M.A. and S.C. performed research; M.A., M.S.T. and K.V.V analyzed data; and M.A., M.S.T. and K.V.V. wrote the paper.

## Additional information

Supplementary information accompanies this paper.

