## Supplementary data for "Effect of ligand sensing on flagellar bundle formation in bacteria"

### MATERIALS AND METHODS

*a.* ***Calculation of bundle geometry parameters*** An *E. coli* with fluorescently stained flagella is shown in Figure S1A. The spread angle of a bundle was measured by forming an angle at a fixed distance of 1 $\mu$m away from the back end of the body such that the complete bundle is contained within this angle. Bundle length was measured as the distance between the polar end of the cell body and the distal end of the flagellar bundle.

*b.* ***Calculation of pitch and radius*** The fluorescently stained cells were cropped from main movie and processed using ImageJ for obtaining helical trajectories. Z-projection of maximum intensity was obtained. For each cell, the pitch and radius were calculated by averaging values at different locations as shown in Figure S1B.

*c.* ***Calculation of torque of rotating tethered E. coli cells*** Torque was calculated based on earlier reported equations (Che et.al, 2008). Tethered cells rotates about rotational axis positioned at either few nanometers away from the cell center (Figure S2A) or cell pole (Figure S2B). To calculate the frictional drag coefficient, cell body is considered to be comprised of a large and small semi-ellipsoid as shown in Figure S2 with $f_L$ and $f_S$ values, respectively.

Following equation was used

$$f_L = \frac{8\pi\eta a_L^3/3}{2\left\{ln\left(\frac{2a_L}{b}\right) - 0.5\right\}} \tag{1}$$

$$f_S = \frac{8\pi\eta a_S^3/3}{2\left\{ln\left(\frac{2a_S}{b}\right) - 0.5\right\}} \tag{2}$$

where $\eta$ is the viscosity of motility medium 9.6 x $10^{-4}$ Ns/m$^2$, $a_L$ is the length of major axis of large ellipsoid ($\frac{L}{2}$ + r) and $a_S$ is the length of major axis of small ellipsoid ($\frac{L}{2}$ - r). 'b' is the cell width which is minor axis length for both the semi-ellipsoids.

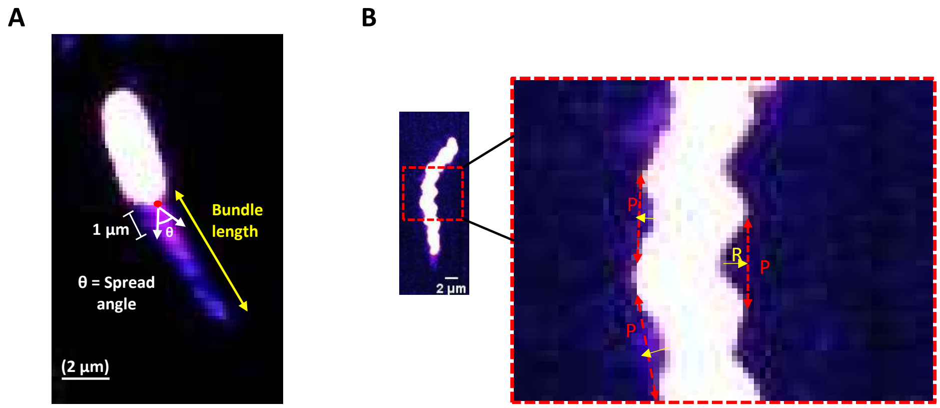


FIG. S1. (A) Illustration for measurement of spread angle and length of flagella. The spread angle is measured at the back end of the cell body and is the minimum angle required to contain all filaments at 1 $\mu$m distance from the back end of cell body. Apparent length is measured from the back of the body to the tip of the bundle (B) Maximum intensity profile obtained for a motile cell in motility buffer to measure the trajectory parameters. 'P' denotes pitch and 'R' radius of helical path. These values are calculated at various points as shown and average values were reported.

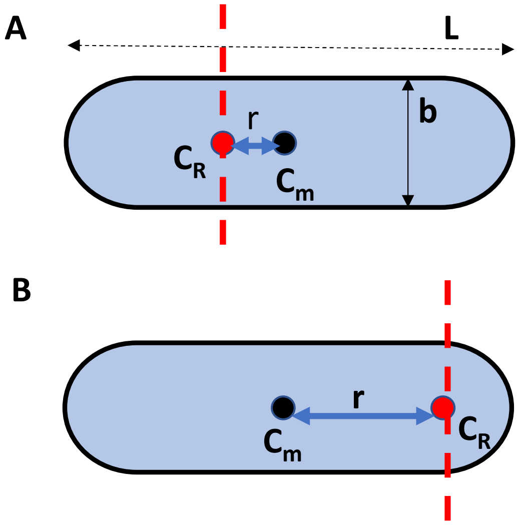


FIG. S2. Schematic of tethered cell describing parameters used for calculating torque. Red line shows the position of rotational axis '$C_R$' which is situated at few nanometers away from centroid '$C_m$' (A) or polar end (B). 'L' is the length of the cell, 'b' is the width and 'r' is the distance between $C_R$ and $C_m$.

Torque (T) is obtained by product of frictional drag coefficient and angular velocity of motor, given by

$T = \omega(f_L + f_S)$ pN.nm, where $\omega$ is angular velocity (rad/s).

### ADDITIONAL RESULTS

*d.* ***Rotational diffusivity in $\Delta cheY$ strain*** $\Delta cheY$ strain has its *cheY* gene deleted and therefore, it is CCW biased resulting in smooth swimming. In MB, the values of $D_r$ are lowest (0.6-0.7 $rad^2$/s) (Figure S3) when compared to WT and $\Delta trg$ strain (Figure 2B-C). Further, the $D_r$ values were reduced in response to glucose (57%) and 2Dg (43%) as compared to MB at 16.5 min of exposure (Figure S3).

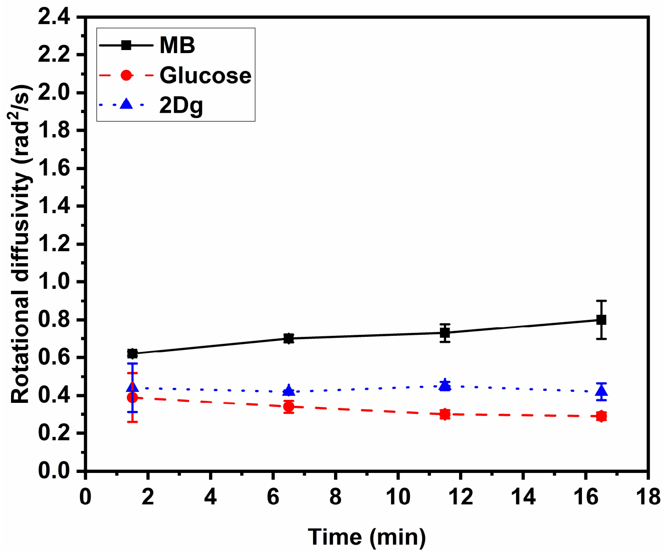


FIG. S3. Time dependent variation in rotational diffusivity for $\Delta cheY$ strain in response to 1000 $\mu M$ glucose and 2Dg as compared to MB. Error bars are standard errors from three independent experiments.

*e.* ***Correlation between run speed and spread angle*** The run speed of $\Delta trg$ cells was plotted against respective spread angle. There was no discernible difference in 2Dg and MB because of absence of Trg receptor (Figure S4A). The cumulative data of run speeds and spread angles obtained from analyzing WT, $\Delta trg$, $\Delta cheY$ in MB, glucose and 2Dg shows that run speed is inversely related to the flagellar spread angle albeit a large spread (Figure

S4B).

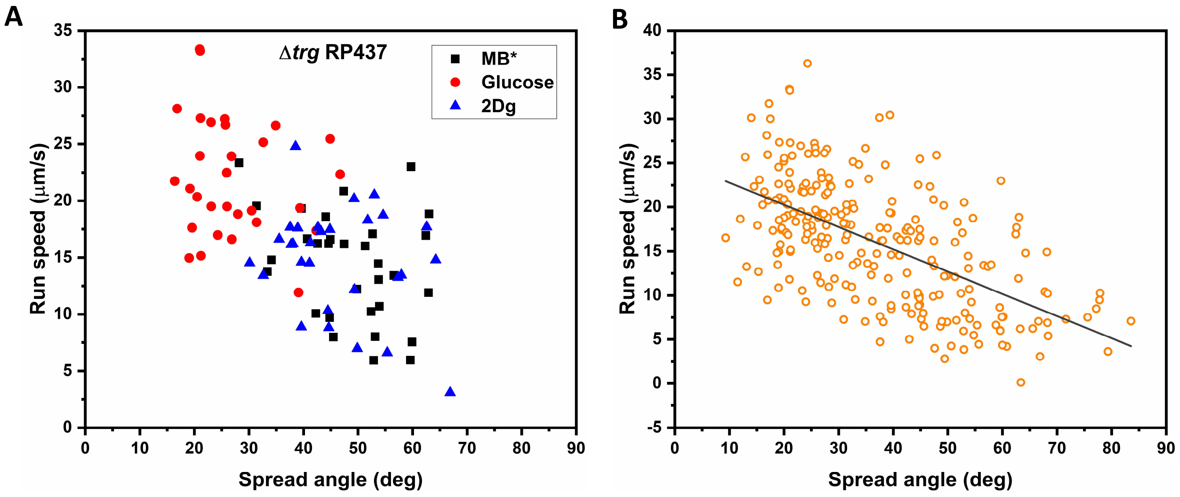


FIG. S4. (A) Run speed and spread angle distribution of Δ*trg* RP437 cells in glucose and 2Dg as compared to motility buffer (B) Cumulative data from WT, Δ*trg*, Δ*cheY* and Δ*ptsI* (≈ 300 cells) in ligands (MB, glucose, 2Dg) shows that run speed is inversely correlated to the spread angle of flagellar bundles
